# Social behavioral profiling by unsupervised deep learning reveals a stimulative effect of dopamine D3 agonists on zebrafish sociality

**DOI:** 10.1101/2021.09.24.461752

**Authors:** Yijie Geng, Randall T. Peterson

## Abstract

It has been a major challenge to systematically evaluate and compare how pharmacological perturbations influence social behavioral outcomes. Although some pharmacological agents are known to alter social behavior, precise description and quantification of such effects have proven difficult. The complexity of brain functions regulating sociality makes it challenging to predict drug effects on social behavior without testing in live animals, and most existing behavioral assays are low-throughput and provide only unidimensional readouts of social function. To achieve richer characterization of drug effects on sociality, we developed a scalable social behavioral assay for zebrafish named ZeChat based on unsupervised deep learning. High-dimensional and dynamic social behavioral phenotypes are automatically classified using this method. By screening a neuroactive compound library, we found that different classes of chemicals evoke distinct patterns of social behavioral fingerprints. By examining these patterns, we discovered that dopamine D3 agonists possess a social stimulative effect on zebrafish. The D3 agonists pramipexole, piribedil, and 7-hydroxy-DPAT-HBr rescued social deficits in a valproic acid-induced zebrafish autism model. The ZeChat platform provides a promising approach for dissecting the pharmacology of social behavior and discovering novel social-modulatory compounds.

## INTRODUCTION

Sociality is broadly conserved across the animal kingdom, facilitating cooperation, reproduction, and protection from predation. In humans, social dysfunction is a hallmark of several neuropsychiatric disorders such as autism, schizophrenia, bipolar disorder, and Williams syndrome, to name a few. In particular, social communication impairment is considered a core symptom of autism. Despite its importance, we lack a comprehensive understanding of how the diverse classes of neuroactive drugs impact social behavior. This is evidenced by the fact that although certain antipsychotics, antidepressants, and stimulants medications are used clinically to help manage some symptoms of autism^1,2^, no treatment is currently available to ameliorate the disease-relevant social deficit.

It has been a major challenge to comprehensively assess and compare how chemicals affect complex behaviors such as sociality. Simple *in vitro* assays cannot effectively model drug effects on whole organisms, especially on brain activity. Rodent models lack sufficient throughput and are cost-prohibitive for a comprehensive examination of the hundreds of neuroactive drugs currently available, limiting their uses to small-scale hypothesis-driven testing. On the other hand, the zebrafish has become an increasingly important model organism for social behavioral research^3^, and recent developments in zebrafish behavioral profiling have demonstrated a promising alternative approach to meeting this challenge. Indeed, multidimensional behavioral profiling in zebrafish has been used to systematically assess thousands of chemicals for effects on motor responses^4,5^, rest/wake behavior^6^, and appetite^7^.

Current methods of social behavioral analysis in zebrafish are mostly limited to quantifying the average measurement of a human-defined simplex trait such as social preference^8^, social orienting^9^, and group cohesion^10^, or a collection of several simplex traits^11^, with limited throughput. Restricted by their unidimensional nature, these measurements often fail to adequately represent the complex and multidimensional nature of social behavior in space and time. To comprehensively assess social behavior for behavioral profiling, we sought to develop an automated method to classify the real-time dynamics of social behavior based solely on information provided by the data, without any human intervention, in a scalable format. To achieve this goal, we adopted an unsupervised deep learning approach: deep learning based on a convolutional autoencoder can automatically extract social-relevant features from a behavioral recording, while unsupervised learning allows for unbiased classification of real-time behavioral phenotypes; both processes were conducted free of human instructions.

Here, we report a fully automated and scalable social behavioral profiling platform named ZeChat. Built on an unsupervised deep learning backbone, ZeChat embeds the high-dimensional and dynamical social behavioral data into a 2-dimensional space and assigns the embedded datapoints to distinct behavioral categories, thus converting a fish’s entire social behavioral recording to a behavioral fingerprint in the form of a numerical vector. Screening 237 known neuroactive compounds using the ZeChat system generated a rich set of social-relevant behavioral phenotypes which enabled unbiased clustering and classification of drug-treated animals. Based on the social behavioral profile compiled from the screen, we discovered a social stimulative effect of dopamine D3 receptor agonists (D3 agonists). Acute exposure to D3 agonists rescued social deficits in a valproic acid-induced zebrafish autism model. Our results demonstrate that multidimensional social behavioral phenotypes can be distilled into simple behavioral fingerprints to classify the effect of psychotropic chemicals on sociality.

## RESULTS

### Rationale and overview of the ZeChat behavioral analysis framework

The ZeChat workflow is summarized in Figure 1a. We probed social interaction in a 2-chamber setup, in which each fish swims freely in a square arena with visual access to its partner fish through a transparent window. In this setup, a fish’s position inside the arena, as well as its posture and movement dynamics, were deemed relevant for social interaction. Inspired by Berman et al.^12^, we sought to describe social behavior as a point moving through a high-dimensional space of positional, postural, and motional features, and to assign segmented subspaces to sub-behaviors. First, a preprocessing step distilled social-relevant information from the recorded images. A convolutional autoencoder then unbiasedly extracted key features from the preprocessed images to a latent vector, which is then projected onto its first 40 principal components. We converted the time series of each principal component to a wavelet spectrogram to incorporate behavioral dynamics into a feature vector. Finally, each feature vector was embedded into a 2-dimensaional map and classified to distinct behavioral categories.

**Figure 1.**
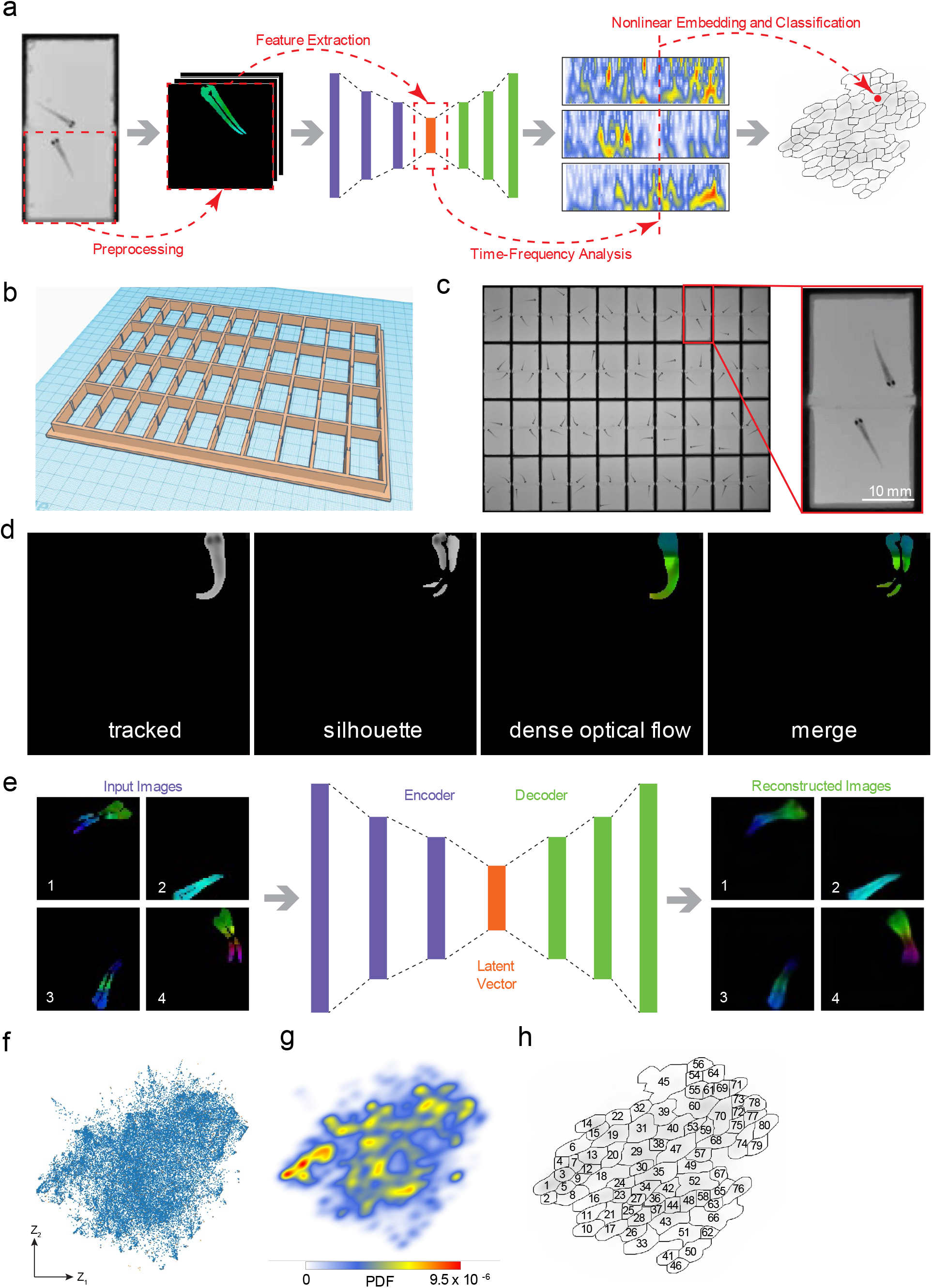
The general framework of ZeChat behavioral analysis. (**a**) To analyze a ZeChat recording, a separate video clip is first generated for each fish by cropping out the ZeChat arena where it is located in. Each cropped video clip is orientated so that the transparent window is always aligned to the top edge of the clip. Each frame is then Preprocessed to preserve positional, postural, and motion related information. The preprocessed images are fed into an autoencoder for Feature Extraction. The main principal components of the extracted feature vector are each converted to a spectrogram by Time-Frequency Analysis. The resulting spectral feature vectors are embedded into a 2-dimensional map and classified to distinct behavioral categories by Nonlinear Embedding and Classification. (**b**) The 3D design of the 40-unit ZeChat testing array. (**c**) A screenshot of ZeChat recording. Also zoom in to show an independent testing unit. (**d**) Intermediate and resulting images of the preprocessing procedure. Fish is first tracked to remove background (tracked). Consecutive tracked frames are subtracted (silhouette). In parallel, the tracked fish is colored by dense optical flow (dense optical flow). Finally, the dense optical flow image is masked by the silhouette to generate a merged image (merge). (**e**) Training the convolutional autoencoder. Preprocessed images (left, Input Images) are fed into the 7-layer convolutional autoencoder (middle) to be reconstructed (right, Reconstructed Images). The Encoder layers are responsible for compressing the input image into a latent representation space in the form of a Latent Vector, which is then used to reconstruct the input image by the Decoder layers. (**f**) Training dataset embedded into a 2D ZeChat map. A reference map containing 3000 datapoints (red) was first embedded using t-SNE. Kernel t-SNE was then used to embed an additional 60,000 datapoints (blue). (**g**) Probability density function (PDF) of ZeChat map containing 10,000 randomly selected datapoints. Generated by convolving the ZeChat map with a Gaussian. (**h**) PDF of the ZeChat map was segmented into 80 distinct behavioral categories by performing a watershed transform.

### Social-relevant information can be extracted via behavioral recording and image preprocessing

The zebrafish becomes socially active at 3 weeks of age^8^ while remaining small in size (~ 1 cm long), enabling us to visualize social interaction in a confined space. To allow easy separation of individual fish for subsequent analysis, pairs of fish were each placed in a separate 2 cm × 2 cm arena and allowed to interact only through a transparent window (Supplementary Fig. 1a; Supplementary Video 1). A custom-built high-throughput imaging platform was used to record 40 pairs of fish simultaneously with sufficient spatiotemporal resolution to capture dynamic changes of the fish’s postures and positions (Fig. 1b-c & Supplementary Video 2). Sexual dimorphism is not readily apparent at this stage, so fish were paired without sex distinction.

For image preprocessing, images of each arena were cropped with the transparent window always in the upright position to preserve fish’s positional information. Each fish was first tracked to be isolated from the background (Fig. 1d & Supplementary Video 3: tracked). Consecutive frames were subtracted to show postural changes between consecutive frames in the resulting silhouette (Fig. 1d & Supplementary Video 3: silhouette). In parallel, we colored each fish based on its instantaneous direction and velocity of movement calculated by dense optical flow^13^ (Fig. 1d & Supplementary Video 3: dense optical flow). Finally, the dense optical flow image was masked by its corresponding silhouette to generate a merged image (Fig. 1c & Supplementary Video 3: merge; Supplementary Fig. 1c).

### Preprocessed images can be transformed to feature vectors by feature extraction and time-frequency analysis

Without any human intervention, convolutional autoencoders can automatically “learn” to extract useful features from input images into a latent vector, which is then used to reconstruct these images. We therefore used this deep learning architecture to extract key features of the preprocessed images into the latent vector for subsequent analyses. As part of the initial setup, we first pre-trained the convolutional autoencoder using a training set of preprocessed images (Fig. 1e & Supplementary Fig. 1d). The resulting latent vectors were projected onto the first 40 principal components by a principal component analysis (PCA), preserving ~ 95% of the total variance. When running the ZeChat analysis, preprocessed images were converted to time series of 40 principal components by the pre-trained autoencoder and PCA models.

Behaviors happen in durations, necessitating time to be taken into consideration to properly interpret information extracted from the behavioral recordings. To embed time-related information into the final feature vector, we adopted the method of applying continuous wavelet transform (CWT) on the time series of each of the 40 principal components to capture oscillations across many timescales^12^. From the 40 resulting spectrograms, 25 amplitudes at each timepoint were concatenated into a single vector of length 40 × 25. Up to this point, each original recorded frame was converted to a single 1,000-dimensional feature vector (Supplementary Fig. 2).

### Feature vectors are assigned to behavioral categories by nonlinear embedding and classification

Finally, we adopted a method developed by Berman et al.^12^, with modifications, to assign feature vectors to behavioral categories through nonlinear embedding and classification. The high dimensional feature vectors were embedded to a 2-dimensional space by nonlinear dimensionality reduction using *t*-distributed stochastic neighbor embedding (*t*-SNE)^14^. Due to computational limitations, we first embedded a small subset of randomly sampled feature vectors to create a reference map. Because *t*-SNE is non-parametric, we applied a parametric variant of *t*-SNE named kernel *t*-SNE^15^ to embed additional datapoints onto the reference map. We named the resulting 2-dimensional behavioral space ZeChat map (Fig. 1f).

Calculating the probability density function (PDF) of ZeChat map identified regions with high datapoint density as local maxima (Fig. 1g), marking the locations of potential behavioral categories^12^. We segmented ZeChat map into 80 regions based on locations of the local maxima using a watershed transform algorithm, allowing each original recorded frame – now embedded as a datapoint in ZeChat map – to be assigned to a particular behavioral category (Fig. 1h).

### The pause-move dynamic of ZeChat map

We made videos to help visualize how a fish’s real-time behavioral changes translate to datapoint trajectories on the ZeChat map (Supplementary Video 4). We found that the trajectory of the 2-dimensional embedding alternates between sustained pauses within certain regions of the map and rapid movements from one region to a distant region on the map. Plotting the velocity of the trajectory revealed a “pause-move” dynamic (Fig. 2a). The low-velocity points were localized in distinguishable peaks that often overlapped with the ZeChat map’s local maxima (Fig. 2b & 1g). In contrast, the high-velocity points were more uniformly distributed (Fig. 2b). This result supports the idea that the social-relevant behavioral changes can be represented by a course through a high-dimensional space of postural, motional, and positional features in which the course halts at locations that correspond to discrete sub-behaviors^12^.

**Figure 2.**
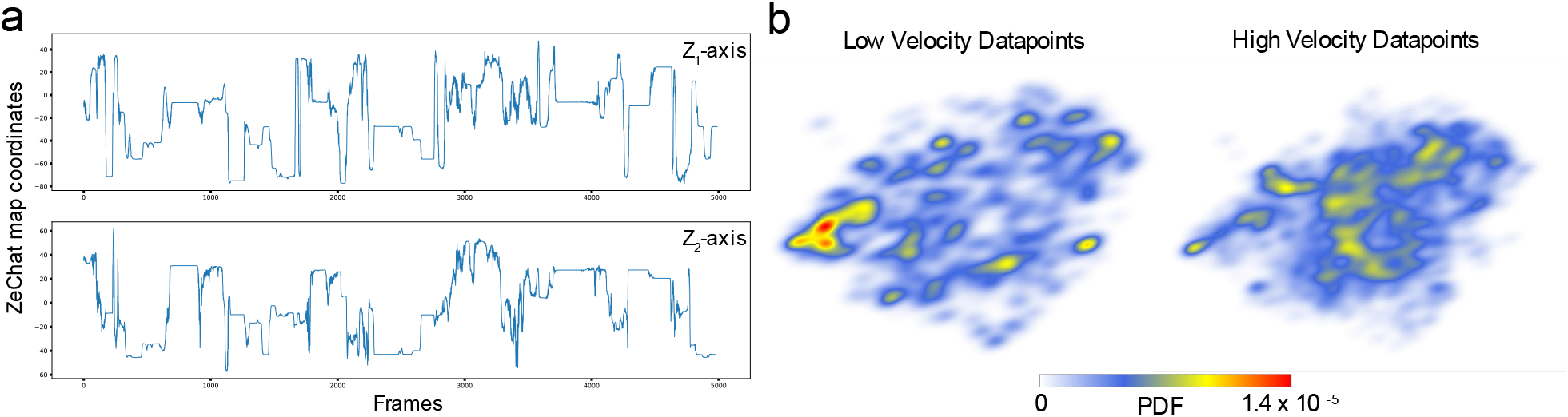
The pause-move dynamic of ZeChat map. (**a**) A typical datapoint trajectory in the Z_1_ and Z_2_ axes of ZeChat map. Showing a pause-move dynamic. (**b**) PDF maps of low velocity (< 1) and high velocity (≥ 1) datapoints. Local maxima positions of the low velocity PDF map closely match local maxima positions in Fig. 2g, whereas the high velocity datapoints showed a more uniform distribution pattern.

### Neuroactive compound screening reveals diverse social behavioral responses

To systematically assess how neuroactive compounds modulate social behavior, we conducted a screen of 237 compounds including modulators of the dopamine, serotonin, and opioid-related pathways. These pathways were selected because they have been implicating in influencing social behavior^16–18^. Briefly, 3-week-old juvenile fish were treated with compounds by bath exposure for 1-3 hours prior to ZeChat recording. Ten fish were treated with each compound, and fish treated with the same compound were paired with each other for ZeChat recording (Fig. 3a). A set of DMSO control fish was included in every recording.

**Figure 3.**
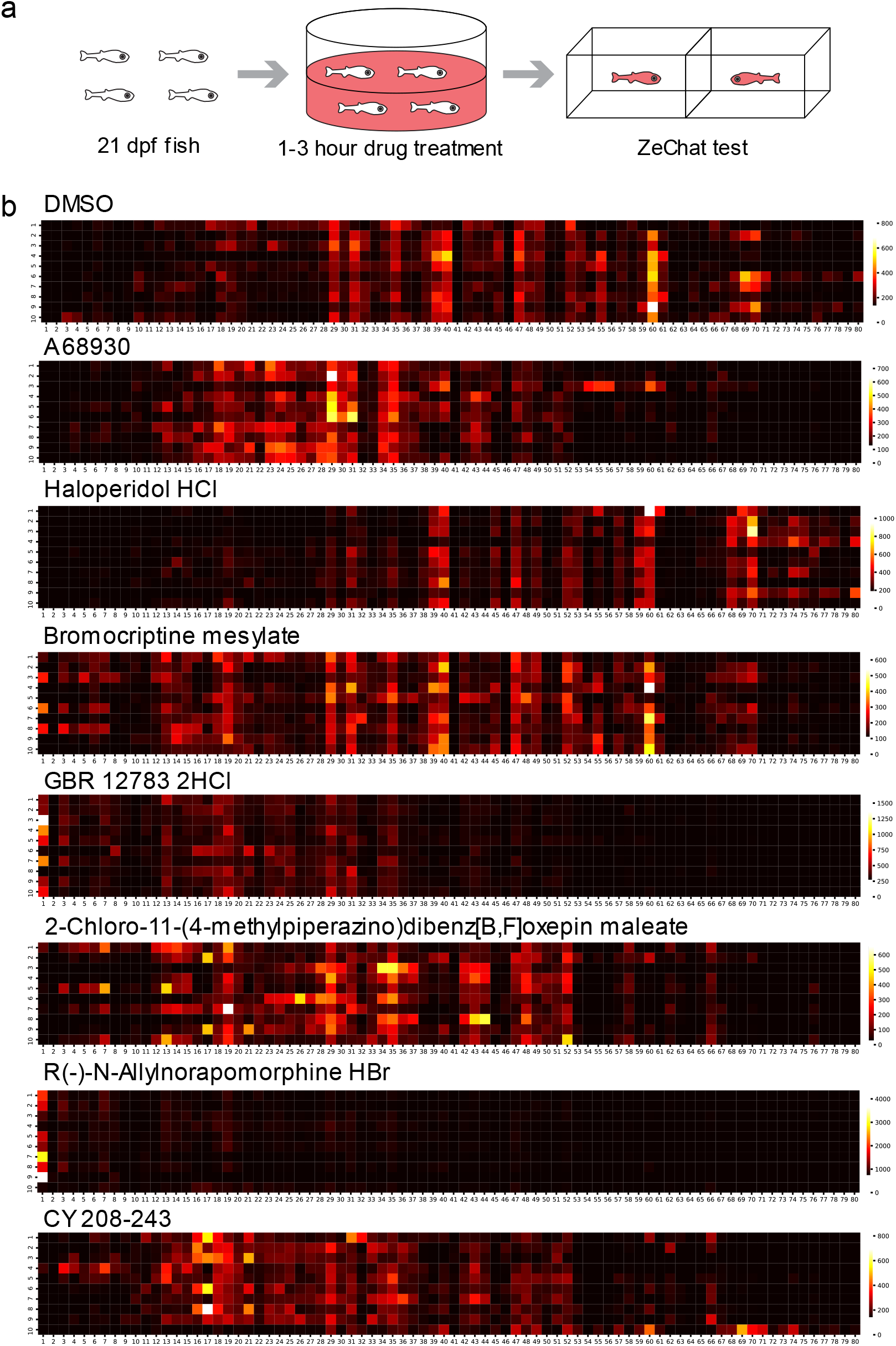
Neuroactive compounds produce highly reproducible behavioral fingerprints. (**a**) A schematic of the screening procedure. (**b**) Behavioral fingerprints of individual fish treated by different chemicals. Each row represents the behavior fingerprint of an individual fish. Each square represents the total number of times a fish is assigned to a behavioral category. Horizontal axis: the 80 behavioral categories. Color bar: cumulated number of times a fish is assigned to a behavioral category.

Counting the number of times a fish’s behavior is classified to each behavioral category generated a behavioral fingerprint in the form of an 80-dimensional numerical vector. Fish treated with the same compound showed highly similar behavioral fingerprints (Fig. 3b), suggesting that the behavioral fingerprints produced by a given compound are consistent across multiple individual animals. To consolidate data, we combined the behavioral fingerprints of fish treated with the same compound by keeping the median value of each behavioral category. All 237 consolidated behavioral fingerprints plus DMSO controls were normalized, and the medians of DMSO controls were subtracted from all samples to help visualize changes in behavioral fingerprints compared to wild type behavior.

Hierarchical clustering reveals a diversity of behavioral responses (Fig. 4 & Supplementary Fig. 3). We found that compounds belonging to the same functional class consistently evoked highly similar behavioral fingerprints (Fig. 5a and Supplementary Fig. 4 & 5). To compare the typical behavioral fingerprints of major drug classes, we calculated the median value of each behavioral category for all behavioral fingerprints elicited by functionally similar molecules. Only drug classes with no fewer than 3 compounds tested in the screen were included in this analysis. Hierarchical clustering of the resulting behavioral fingerprints again revealed distinct behavioral phenotypes (Fig. 5b). Remarkably, compounds targeting the 3 major neurotransmitter pathways, e.g., the serotonin, dopamine, and opioid pathways, were naturally separated by hierarchical clustering (Fig. 5b: functional classes of drugs are color coded to distinguish the 3 major pathways).

**Figure 4.**
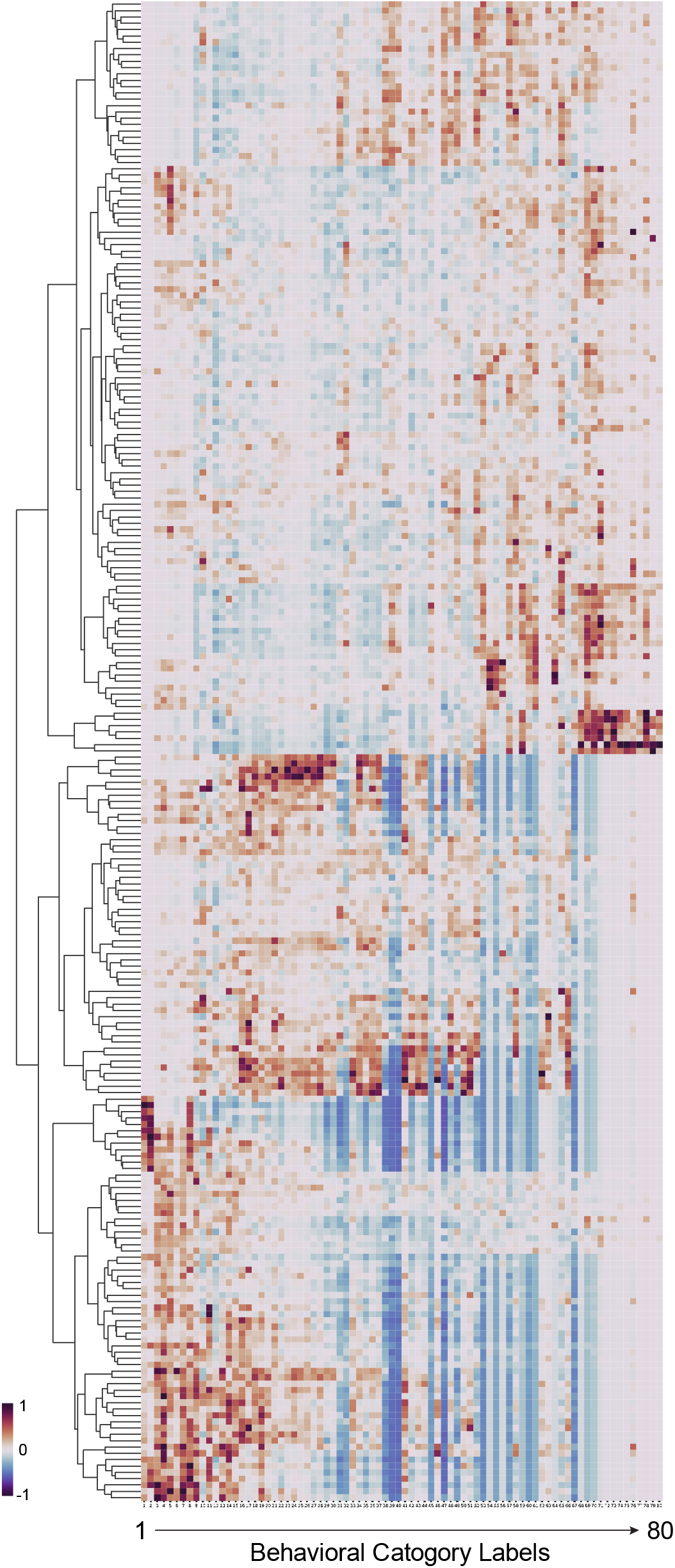
Hierarchical clustering reveals distinct drug-induced behavioral responses. Hierarchical clustering of behavioral fingerprints generated by the screen. Each behavioral fingerprint (row) represents the median value of the individual fingerprints of all fish (n≤10 per treatment) treated by the same compound. The behavioral fingerprints are normalized for each behavioral category and subtracted by the median DMSO fingerprint. Horizontal axis labels the 80 behavioral categories.

**Figure 5.**
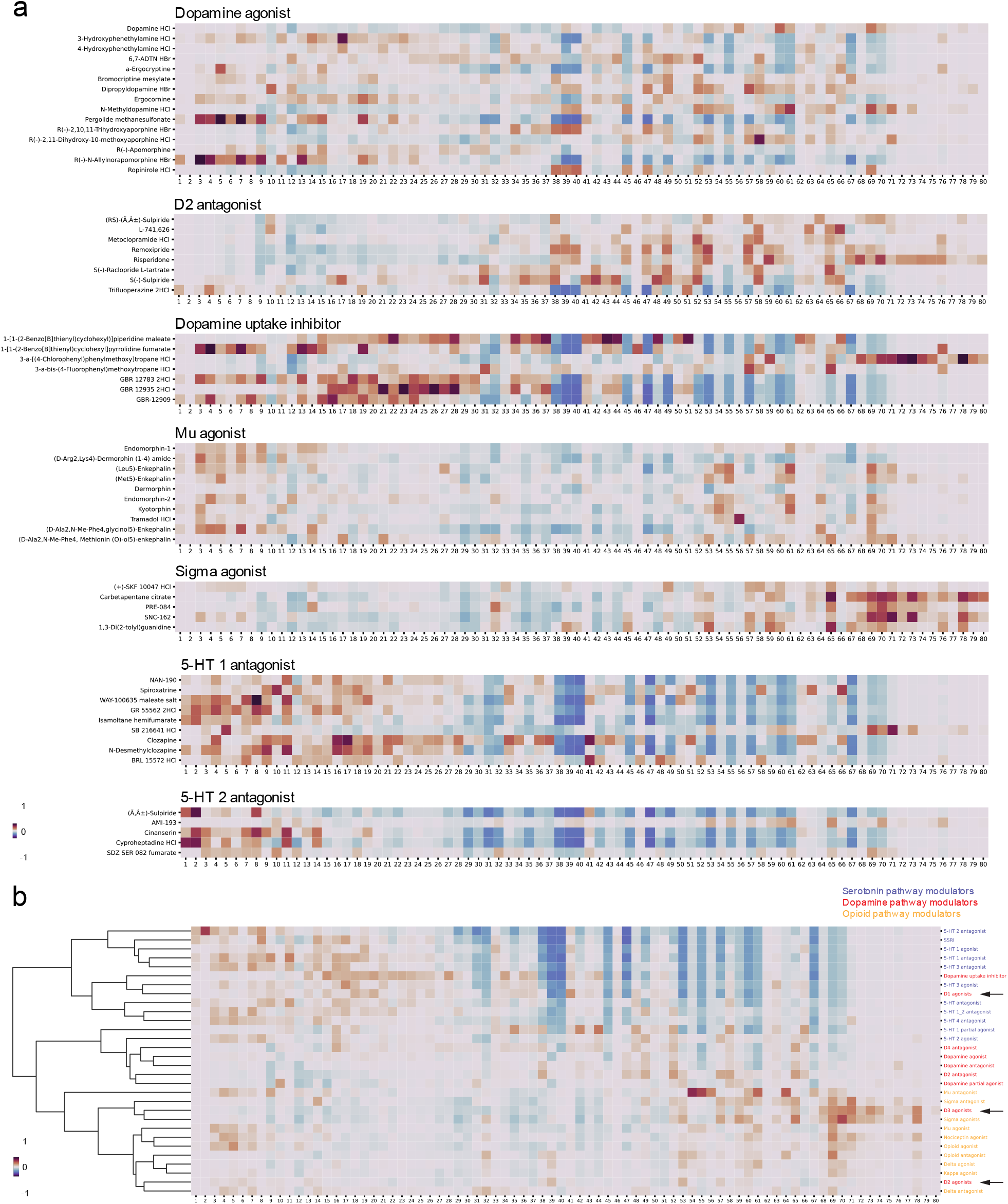
Functionally similar molecules evoke similar behavioral responses. (**a**) Neuroactive compounds with similar annotated functions elicit similar behavioral fingerprints. (**b**) Behavioral fingerprints of functionally similar molecules are consolidated to a single behavioral fingerprint by calculating the median value of each behavioral category, and the resulting behavior fingerprints are hierarchically clustered. Only groups of drugs containing no less than 3 compounds sharing the same annotated function are included in the analysis. The group labels are colored by the targeted pathway. Black arrows point to behavioral fingerprints of dopamine D1, D2, and D3 receptor agonists, respectively.

### Dopamine D3 receptor agonists rescue social deficits in a VPA-induced autism model

Surprisingly, we noticed that the dopamine D1, D2, and D3 receptor agonists were clustered well apart from each other (Fig. 5b: black arrows), suggesting that selected activations of the dopamine D1, D2, and D3 receptor-related neuronal circuits elicited distinct social behavioral phenotypes. The five D3 agonists tested in the 237-compound screen generated highly similar behavioral fingerprints sharing a unique pattern in which strong signals are observed in the higher-number behavioral categories (Fig. 6a-b). In contrast, the D1 and D2 agonists elicited very different behavioral fingerprints with no enrichment in these higher-number behavioral categories (Fig. 6a). By examining raw behavioral recordings, we noticed that the D3-agonist-treated fish tend to spend a significant amount of time swimming intensively while pressing against the transparent window. Compared to wild type animals, these fish demonstrated persistent and strong high-frequency tail beats, fast swim velocity, and quick and frequent turns; they also rarely retreated from proximity to the transparent window (Supplementary Video 5 and Fig. 6c). We hypothesized that these D3 agonist-associated behaviors may signify enhanced sociality.

**Figure 6.**
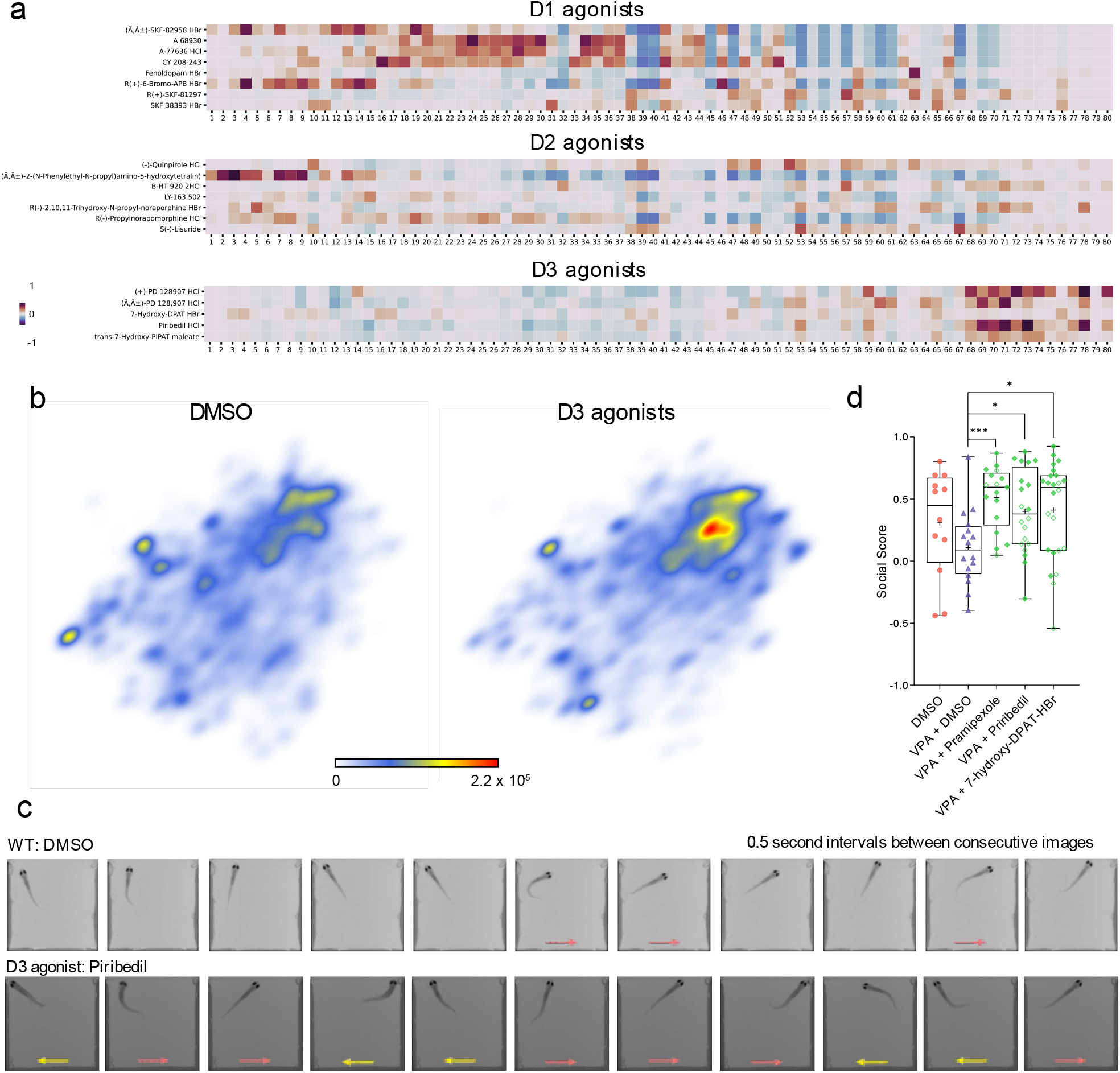
Dopamine D3 agonists rescue social deficits in VPA-treated fish. (**a**) Comparing the behavioral fingerprints of D1, D2, and D3 agonists. The behavioral fingerprints are normalized and subtracted by the median DMSO fingerprint. (**b**) PDF maps of DMSO-treated fish (n=356) and fish treated by D3 agonists (n=49). (**c**) Series of images taken at 0.5 second intervals reveal different swim dynamics between wild type treated by DMSO and fish treated by the D3 agonist piribedil (10 μM). Arrows in red and yellow point to fish’s direction of movement in the current frame. (**d**) Boxplot showing social preference (social score) of DMSO-treated fish (n=12) or VPA-treated fish acutely exposed to DMSO (n=16) or 10 μM D3 agonist including pramipexole (n=17), piribedil (n=20), and 7-hydroxy-DPAT-HBr (n=24) for 1 hour before social preference test. In each boxplot, box encloses data points from the 25^th^ percentile to the 75^th^ percentile, the horizontal line and cross mark the median and the mean, the lines above and below the box reach datapoints with the maximum and minimum values. *: *p*<0.05, ***: *p*<0.001.

We attempted to validate the hypothesized social stimulative property of D3 agonists in a zebrafish autism model with a social deficit phenotype. Embryonic exposure to valproic acid (VPA) is an established model of autism in rodents^19^ and zebrafish^20^. Using a simple zebrafish social preference assay^21^, we observed a clear social deficit phenotype in VPA-treated zebrafish (Supplementary Fig. 6a). To test the effect of D3 agonists against social deficits, we acquired 3 structurally diverse D3 agonists, pramipexole, piribedil, and 7-hydroxy-DPAT-HBr (Supplementary Fig. 6b). Both pramipexole and piribedil are FDA-approved antiparkinsonian agents. We found that exposure to D3 agonists for 1 hour by simple submersion prior to the social preference assay effectively rescued the social deficit in the VPA-treated fish (Fig. 6d).

## DISCUSSION

ZeChat is a deep learning-based behavioral assessment tool enabling scalable and low-cost zebrafish behavioral profiling to characterize changes in sociality. The *in vivo* ZeChat platform combines advantages of *in vitro* and rodent models, enabling scalable testing with high behavioral resolution. Compared to previous zebrafish behavioral profiling methods, the ZeChat analysis method specifically processes and analyzes social behavior-relevant information, linking known neuroactive drugs with complex but distinct social behavioral outcomes.

Apart from unsupervised machine learning, alternative approaches are available for improving the resolution of social behavioral analysis, but not without drawbacks. For example, supervised machine learning methods have been widely adopted to analyze social interactions in fruit fly^22,23^, zebrafish^24^, and mouse^25^. However, this method still relies on human interpretation of animal behavior to classify and assign behavior and is likely unable to fully reveal the complexity and subtilty of social behavior. Another approach uses predefined measurement criteria to mathematically model and classify social interaction^26,27^, which reduces human biases in the analysis, but the quality of its outcome is highly dependent upon the validity of the model. In comparison, unsupervised methods have successfully revealed stereotypic behavioral motifs in individual animals of C. elegans^28–34^, fruit fly^12,35–39^, zebrafish^40–42^, and mouse^43,44^, as well as paired interactions in fruit fly^45,46^, without any human interventions or *a priori* assumptions, providing a viable approach for our purpose.

However, all these approaches still rely on manual selection of features for data preprocessing, which requires strong domain knowledge in the behaving animal. These prerequisites are not always met, especially when faced with complex problems such as analyzing subtle behavioral changes in a video or analyzing sequences of behaviors, as it is difficult for a human observer to exhaustively extract useful features from an image or a sequence of images. Deep learning methods, on the other hand, can automatically learn to extract abstract features from images. As behavioral recordings are sequences of images, the potential benefit for applying deep learning to process these data is apparent. In fact, several recent studies have successfully utilized deep learning to facilitate individual animal identification^47^, tracking^48^, and movement prediction^49^ in zebrafish, paving the way for its application in ZeChat.

In alignment with our findings, the D3 receptor has been previously implicated in social behavioral regulation. In humans, pramipexole alleviates social anxiety in selective serotonin reuptake inhibitor (SSRI)-treated patients^50^. In rodents, two D3 agonists 7-OH-DPAT and PD 128907 were reported to cause a variety of complex alterations in social behavior^51,52^. Further investigations are needed to validate these findings in rodents using other D3 agonists and under different test conditions, drug doses, and genetic backgrounds of the animals, but the results in zebrafish, rats, and humans all point to an important role of D3 receptors in modulating social behavior. In addition, because both pramipexole and piribedil are FDA-approved antiparkinsonian agents, it may be worthwhile examining their impact on the social behavior of patients receiving these drugs.

Future studies using the ZeChat platform may expand to screening other neuroactive compounds, compounds with no known neuroactivity, and uncharacterized compounds, in the hope of identifying additional phenotypes and drug classes with social-modulatory properties. The characteristic behavioral fingerprint of the D3 agonists may be used to discover novel compounds with similar behavioral effects. In addition to wild type fish, fish carrying mutations relevant to human psychiatric disorders can also be assayed, and their behavioral fingerprints compared to the neuroactive compound clustergram to associate genetic mutations with perturbations of neuronal pathways. As demonstrated by Hoffman et al.^53^, small molecules evoking an anti-correlated behavioral fingerprint may ameliorate social deficits in the mutant fish. Hence, by providing a rapid, high-resolution means of characterizing and categorizing zebrafish with altered social behaviors, ZeChat represents a useful tool for investigating the role of genes and pharmacological agents in modulating complex social behaviors.

## MATERIALS AND METHODS

### The ZeChat imaging system setup

The basic unit of this system is a 10 mm deep, 20 mm wide, and 41.5 mm long (internal dimension) rectangular chamber with 2 mm thick walls. A 10 × 4 array consist of 40 independent testing units was 3D printed using white PLA at 100% infill. The printed test arena was glued onto a 3/16” thick white translucent (43% light transmission) acrylic sheet (US Plastic) using a silicone sealer (Marineland). Each unit was then divided into two square-shaped compartments by inserting a 1.5 mm thick transparent acrylic window – precision cut to 10 mm x 41 mm pieces using a laser cutter – into 0.5 mm deep printed slots located in the middle of each unit on the side of the 41.5 mm wall and fastened using the silicone sealer.

The key component of the imaging system is a 322 mm diameter bi-telecentric lens (Opto Engineering) with an IR (850 nm) bandpass filter (Opto Engineering). A telecentric lens only allows passing of light that is parallel to the optical axis, thus avoiding parallax error in imaging, and enables all test units – being located either in the middle or close to the edge of the field of view – to be imaged without distortion. Videos were taken at 50 frames per second (fps) by a 75 FPS Blackfly S Mono 5.0 MP USB3 Vision camera (PointGrey) with a resolution of 2448 × 2048. The tail beat frequency (TBF) for adult zebrafish is ~ 20 Hz^54^, therefore images taken at 50 Hz by the camera should adequately sample motion-relevant features based on the Nyquist–Shannon sampling theorem. The imaging platform was back-illuminated with an infrared (850 nm) LED array (EnvironmentalLights) to provide light for video recording. The infrared LED array was positioned on top of a heat sink (H S Marston). The imaging platform was also illuminated from two opposing sides using white LED arrays (EnvironmentalLights) to provide ambient light for the test subjects. Structural supports and enclosure were custom built using parts purchased from Thorlabs, McMaster Carr, and US Plastic.

### ZeChat test

Test subjects were individually placed into each unit – one on each side of the transparent window – using a transfer pipette with its tip cut off. Their visual access to each other was temporarily blocked by a 3-D printed nontransprent comb-like structure (Supplementary Fig. 1b) prior to each recording session. Once all test subjects were placed into test arenas, the entire test apparatus was transferred into the imaging station and the combs were removed to allow visual access between each pair of fish.

The 2-compartment social interaction setup allows the behavior of each fish to be recorded and analyzed independently without having to go through complex and often computationally demanding and time-consuming tracking procedures to separate each fish. Videos were streamed and recorded using the software Bonsai^55^. A 10 min test session was video recorded for each test. To give fish an acclimation period at the beginning of each test and to take into consideration that the effects of some of the drugs tend to wear off quickly, only the 5 min video segment between 2.5 min and 7.5 min was used for subsequent analyses. All subsequent data processing and analyses were conducted in Python using packages including OpenCV, scikit-learn, Keras, PyWavelets, and imutils.

### Data preprocessing

For data preprocessing, individual fish were first separated from the background using the K-nearest neighbors method^56^. A separate video segment was cropped out for each fish which contains a recording of the entire square compartment where the fish is located. Because the relative position of a fish to its compartment is relevant to social interaction dynamics, each compartment was analyzed as a whole. And because each compartment is polarized, with only one of the four sides being transparent to another fish, for each pair of compartments, the video containing fish in the “top” compartment is flipped vertically by rotating 180 degrees to match the orientation of video recording the “bottom” compartment, so that the side of the compartment facing the transparent widow always faces upward in each video.

To capture changes in each fish’s posture between consecutive frames, we subtracted every current frame from its previous frame. The resulting images were binary-thresholded to generate silhouette-like masks. In parallel, we calculated each fish’s direction of movement between consecutive frames using the Franeback Method of dense optical flow^13^ and used this information to color the fish; motionless fish appear dark after applying this method, thus restricting our analysis to fish in motion. Finally, we applied the mask acquired by subtracting consecutive frames to the dense optical flow image so that the image colored by dense optical flow is cropped by the subtracted silhouette-like mask.

### Training the convolutional autoencoder and feature extraction

The architecture of the convolutional autoencoder consists of three encoding layers each containing 64, 32, and 16 filters, and three decoding layers each containing 16, 32, and 64 filters. We used a training set of preprocessed images to pre-train the convolutional autoencoder. The preprocessed images with a dimension of 220 pixels × 220 pixels were first resized to 56 pixels × 56 pixels to reduce computational requirements. Because a wild type fish typically spends most of the time interacting with its paired fish by staying close to the transparent window, causing the position of the fish in input images to be highly polarized, we enriched the training dataset by rotating each resized image by 90°, 180°, and 270° to generate input images with more postural and positional variations.

The autoencoder forces input images to pass through a “bottleneck” before reconstruction. The bottleneck, or the latent representation space, has a dimension of 784. We then applied principal component analysis (PCA) to this 784-dimensional feature vector and extracted 40 principal components which preserved ~ 95% of total variance.

### Time-frequency analysis of feature dynamics

Calculating the 40 principal components for each video frame yields 40 timeseries for each video. Each timeseries was then expanded into a spectrogram by applying the Continuous Wavelet Transform (CWT). The Morlet wavelet was used as the mother wavelet and 25 scales were chosen to match frequencies spanning from 0.38 Hz to 5 Hz, with the range of frequencies empirically determined to preserve the strongest signals. The time-frequency representation augments the instantaneous representation by capturing oscillations across many timescales. The spectral amplitudes of each time point were then concatenated into a vector of length 40 × 25, giving rise to a 1,000-dimensional representation for each original video frame. Each 1,000-dimensional vector was normalized to having a sum of 1 in order to treat each vector as a probability distribution for subsequent calculation.

### Nonlinear embedding and segmentation

We then performed nonlinear dimensionality reduction on these high dimensional vectors using the popular nonlinear manifold embedding algorithm *t*-distributed stochastic neighbor embedding (*t*-SNE)^14^. We randomly selected and embedded 3,000 feature vectors from 60 fish to generate a reference map. The *t*-SNE algorithm is non-parametric. Therefore, additional datapoints were embedded onto the reference map using a parametric kernel *t*-SNE^15^ method to form the ZeChat map. As the feature vectors are normalized and treated as probability distributions, we calculated the Jensen–Shannon distance (the square root of the Jensen–Shannon divergence) between each pair of vectors as a distance metric for both *t*-SNE and kernel *t*-SNE. We chose the Jensen–Shannon distance as a metric for calculating distances due to it being symmetric and bounded by 0 and 1 which avoids the generation of infinite values.

We calculated the probability density function (PDF) of this map by convolving with a Gaussian kernel. Due to computational limitations, this calculation was conducted using a ZeChat map containing 10,000 randomly selected datapoints. The resulting probability density map was then inverted to turn local maxima into “valleys”. The “ridges” between valleys were detected using Laplacian transform. Finally, a watershed transform was applied to mark the borders between each valley to unbiasedly segment the ZeChat map into 80 behavioral categories.

For ZeChat analysis, to reduce computation time, we randomly sampled 5000 frames from each fish for kernel *t*-SNE embedding and subsequent analyses.

### Behavioral fingerprint calculation and hierarchical clustering

Each frame is assigned a watershed region (behavioral category) based on ZeChat map segmentation. For each fish, the total number of frames assigned to each watershed region was counted, giving rise to a behavioral fingerprint in the form of an 80-dimensional vector. Behavioral fingerprints of fish treated by each drug were combined into one fingerprint by calculating the median of each behavioral category. All combined raw behavioral fingerprints were normalized so that the signals of each behavioral category were between 0 and 1. To help visualize the difference in behavioral patterns between drug treatments and DMSO control, we calculated the median of each behavioral category of all DMSO controls to generate a representative fingerprint for DMSO control, and subtracted this fingerprint from all drug treatment samples. Finally, the normalized and DMSO-subtracted fingerprints of each drug treatment were clustered using the clustermap function (metric=‘euclidean’, method=’complete’) of Python’s Seaborn library.

### Zebrafish chemical treatment and screening

For ZeChat testing, 21 dpf zebrafish were collected from nursery tanks. Fish of roughly average size were selected to minimize the effect of size differences. For the screen, 10 fish were picked into a 60 mm petri dish containing 10 ml E3 medium. Compounds were then added to each dish at a final concentration of 10 μM (non-peptide molecules) or 1 μM (endogenous neuropeptides and their analogs). Fish were incubated for 1-3 hours prior to ZeChat testing. Immediately before testing fish in a petri dish, the content of the petri dish was poured through a nylon tea strainer to remove liquid while keeping fish in the tea strainer. The tea strainer was then consecutively dipped into 3 petri dishes containing E3 to wash the residual chemical away from the fish. The fish were then poured into a petri dish containing clean E3 and each individual was transferred into the ZeChat test arena using a plastic transfer pipette for testing.

### Rescue of VPA fish and social preference testing

VPA treatment was conducted by submerging embryos in 1 μM VPA in E3 medium from 0 to 3 dpf. The drug treated embryos were washed at 3 dpf and transferred to petri dishes containing clean E3 medium. At 5-7 dpf, larvae were transferred into nursery tanks and raised to 21 dpf for behavioral testing of social preference using a 3-chamber assay apparatus^21^. For the D3 agonist rescue experiment, 20 VPA-treated fish were picked into a 25 mm deep 10 cm petri dish containing 30 ml E3 medium. Compounds were then added to each dish and fish were incubated for 1 hour. Immediately before testing, fish were washed as described above, and individually placed into the social preference testing arenas for behavioral testing.

### Chemical library and other compounds

All screening compounds were acquired from the Biomol neuroactive compound library (Biomol) which contains a total of 700 neuroactive drugs dissolved in DMSO at a stock concentration of 10 mM or 1 mM (for only a small subset of drugs). Valproic acid was purchased from Sigma-Aldrich. Pramipexole was purchased from Cayman Chemical. Piribedil was purchased from Selleck Chemicals. 7-hydroxy-DPAT-HBr was purchased from Santa Cruz. All individually purchased compounds were dissolved in DMSO. Chemical structures were generated using PubChem Sketcher.

### Zebrafish husbandry

Fertilized eggs (up to 10,000 embryos per day) were collected from group mating of EkkWill strain zebrafish (*Danio rerio*) (EkkWill Waterlife Resources). Embryos were raised in HEPES (10 mM) buffered E3 medium at 28°C, with or without compound treatment, during the first 3 days. At 3 days post fertilization (dpf), chorion debris was removed, and larvae were transferred into petri dishes containing fresh E3 medium. At 5 – 7 dpf, larvae were transferred into nursery tanks and raised at 28°C on a 14/10 hr on/off light cycle.

### Statistical analysis

Graphs were generated using GraphPad Prism or Python using the Matplotlib package. Data were analyzed using the 2-tailed Student’s *t*-test. P values less than 0.05 were considered significant.

### Code availability

Code is available on the GitHub repository at https://github.com/yijie-geng/ZeChat and is archived on Zenodo under DOI: 10.5281/zenodo.5519964.

## AUTHOR CONTRIBUTIONS

Y.G. conceived the study, built the equipment, designed and conducted the experiments, wrote the Python codes, analyzed the data, and wrote the manuscript. R.T.P conceived the study, designed the experiments, interpreted the data, and revised the manuscript.

## ACKNOWLEDGEMENTS

We thank members of our research group for helpful advice. This work was supported by the L. S. Skaggs Presidential Endowed Chair and by the National Institute of Environmental Health Sciences of the National Institutes of Health under Award Number K99ES031050. The content is solely the responsibility of the authors and does not necessarily represent the official views of the National Institutes of Health.

## SUPPLEMENTARY DATA

**Supplementary Figure 1.**
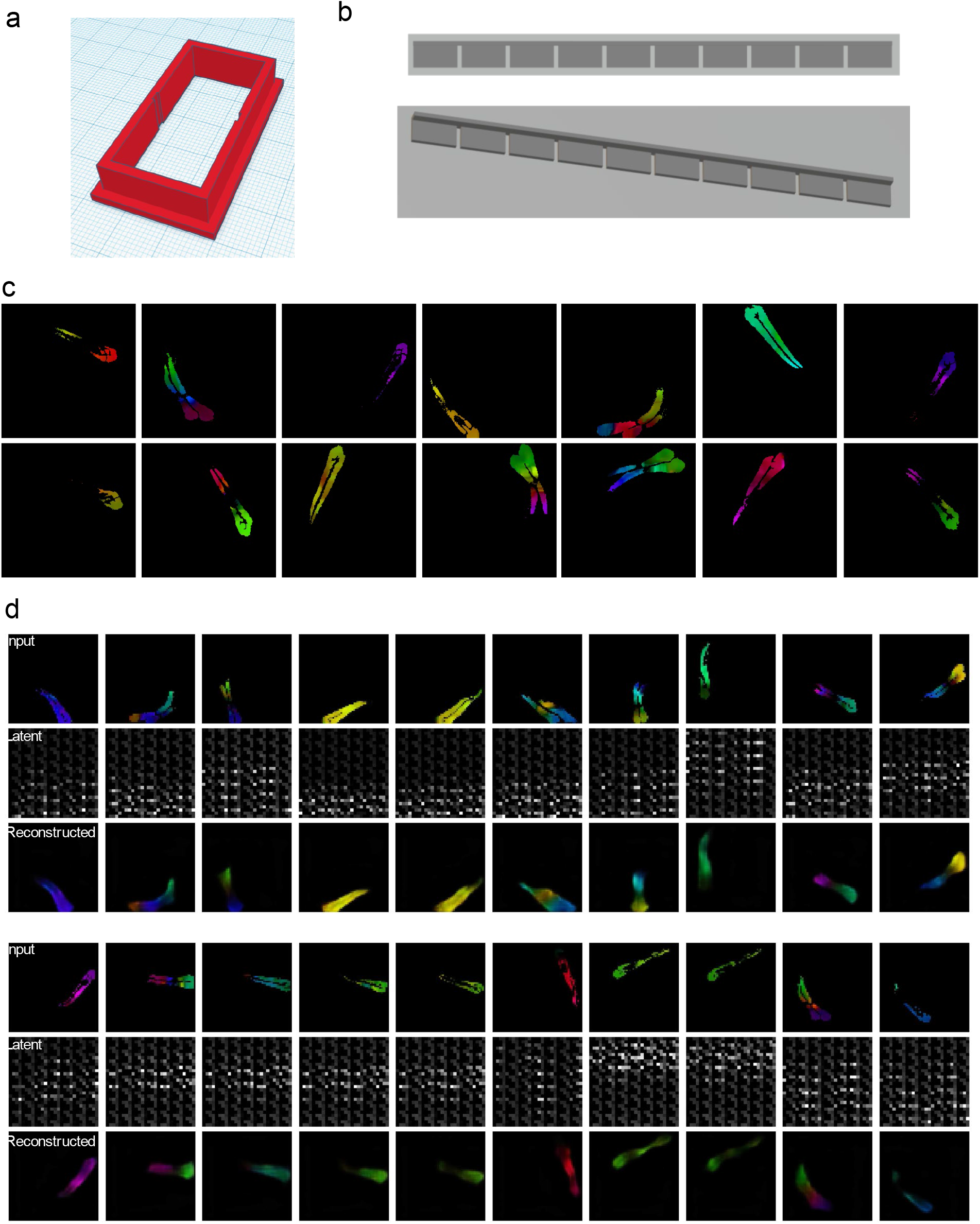
(**a**) The 3D design of one ZeChat unit. (**b**) The 3D design of a comb-like insert for blocking the views of fish before ZeChat test. (**c**) Example preprocessed images. (**d**) Example input images, latent vectors, and reconstructed images.

**Supplementary Figure 2.**
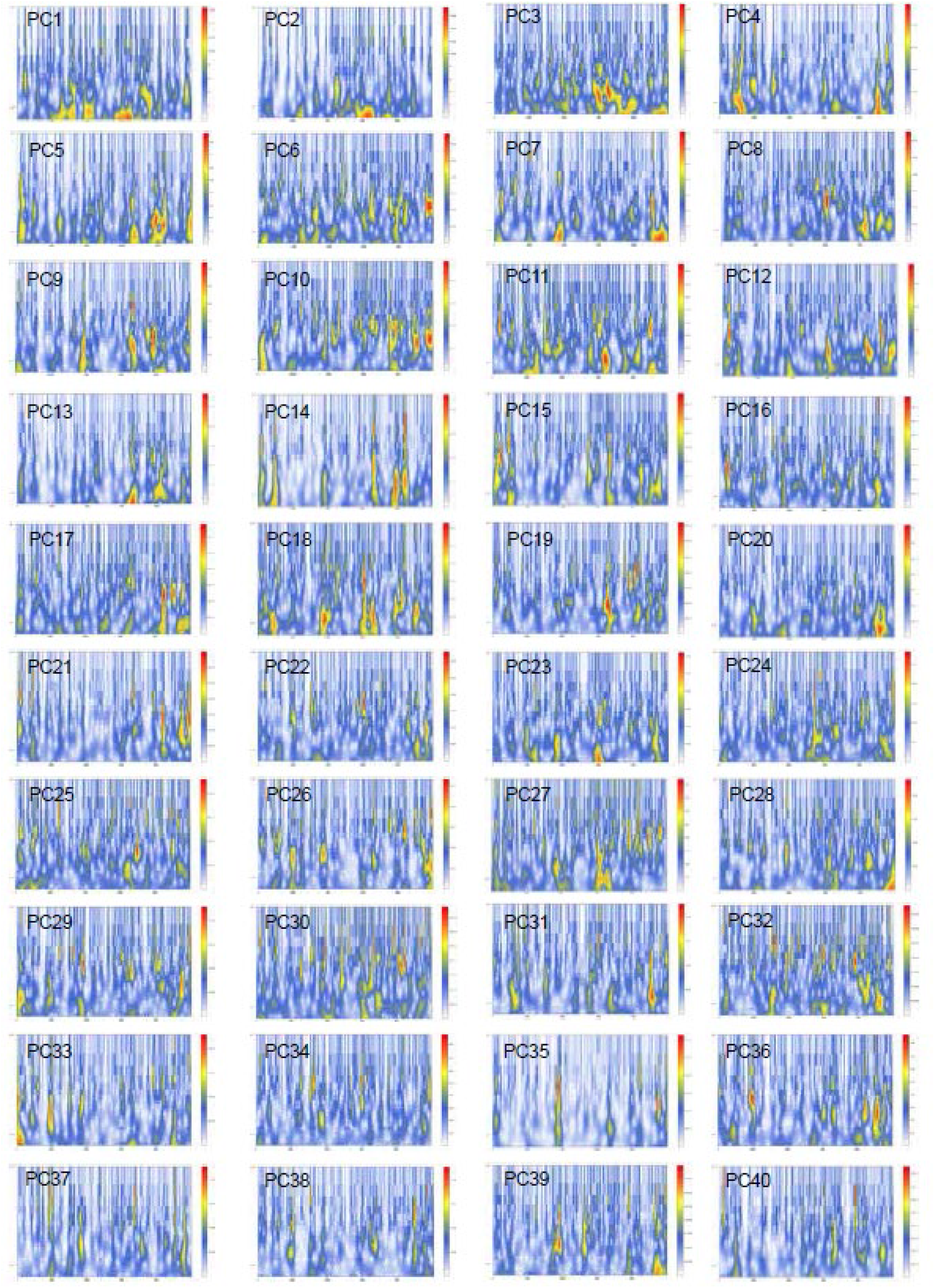
An example of spectrograms generated by time-frequency analysis of 40 principal components of a latent vector. PC1-40: principal components 1-40. Horizontal axis: frames. Vertical axis: frequencies. Color bar: amplitudes.

**Supplementary Figure 3.**
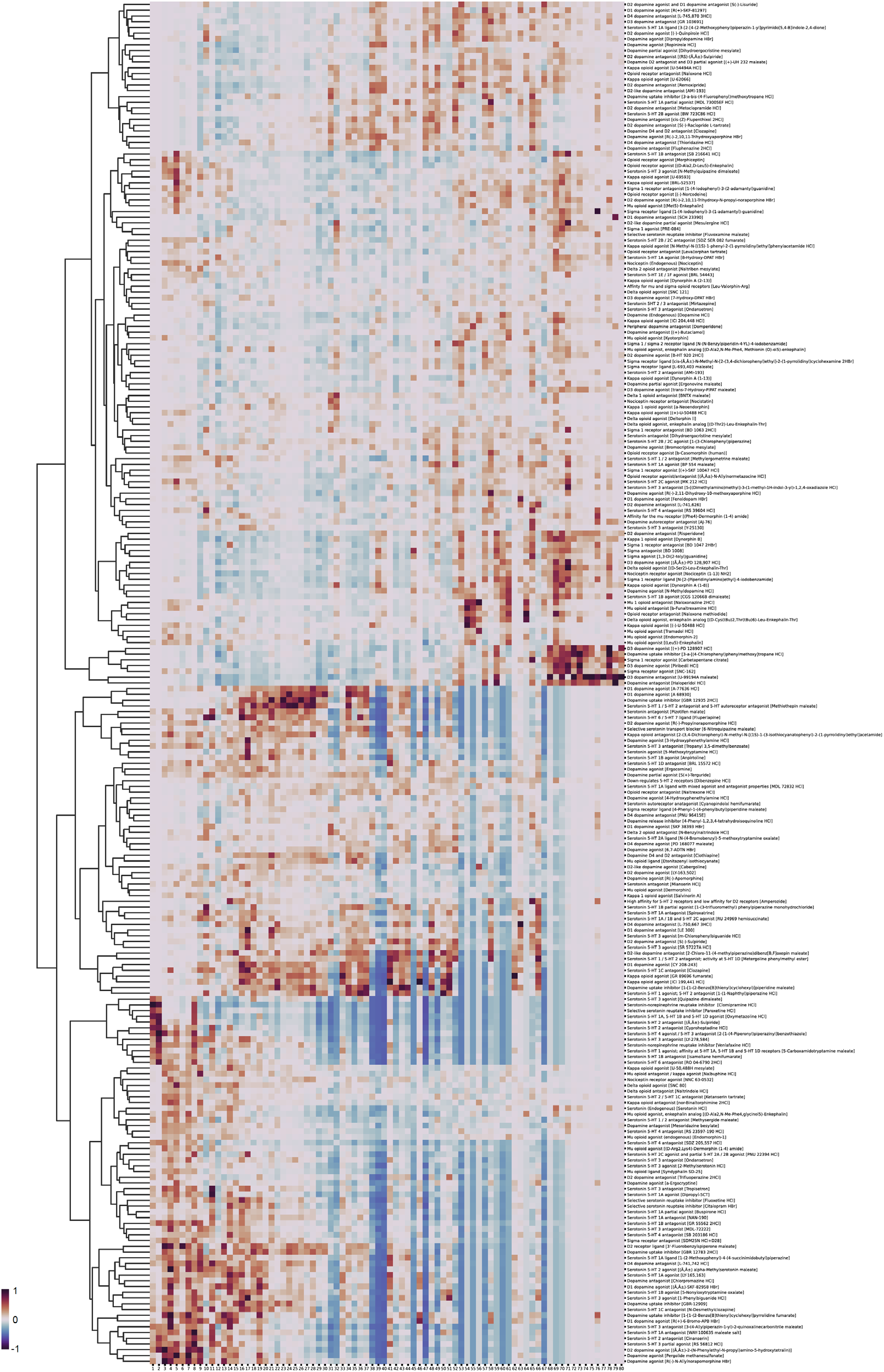
Hierarchical clustering of 237 behavioral fingerprints generated by the screen. The behavioral fingerprints are normalized and subtracted by the median DMSO fingerprint. Labels on the right show: drug classification [drug name].

**Supplementary Figure 4.**
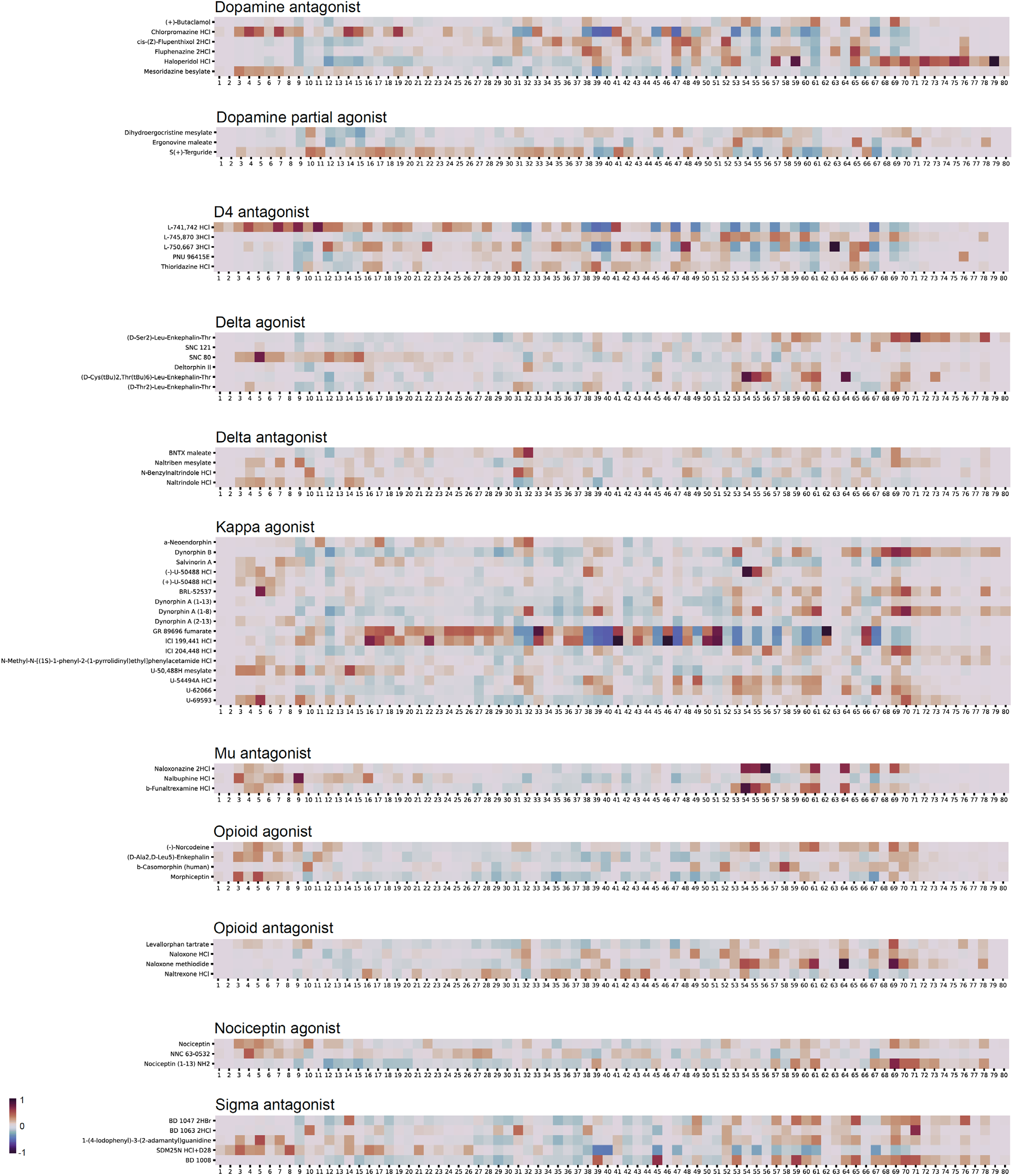
Behavioral fingerprints of dopamine pathway and opioid pathway modulators, grouped by drug effects.

**Supplementary Figure 5.**
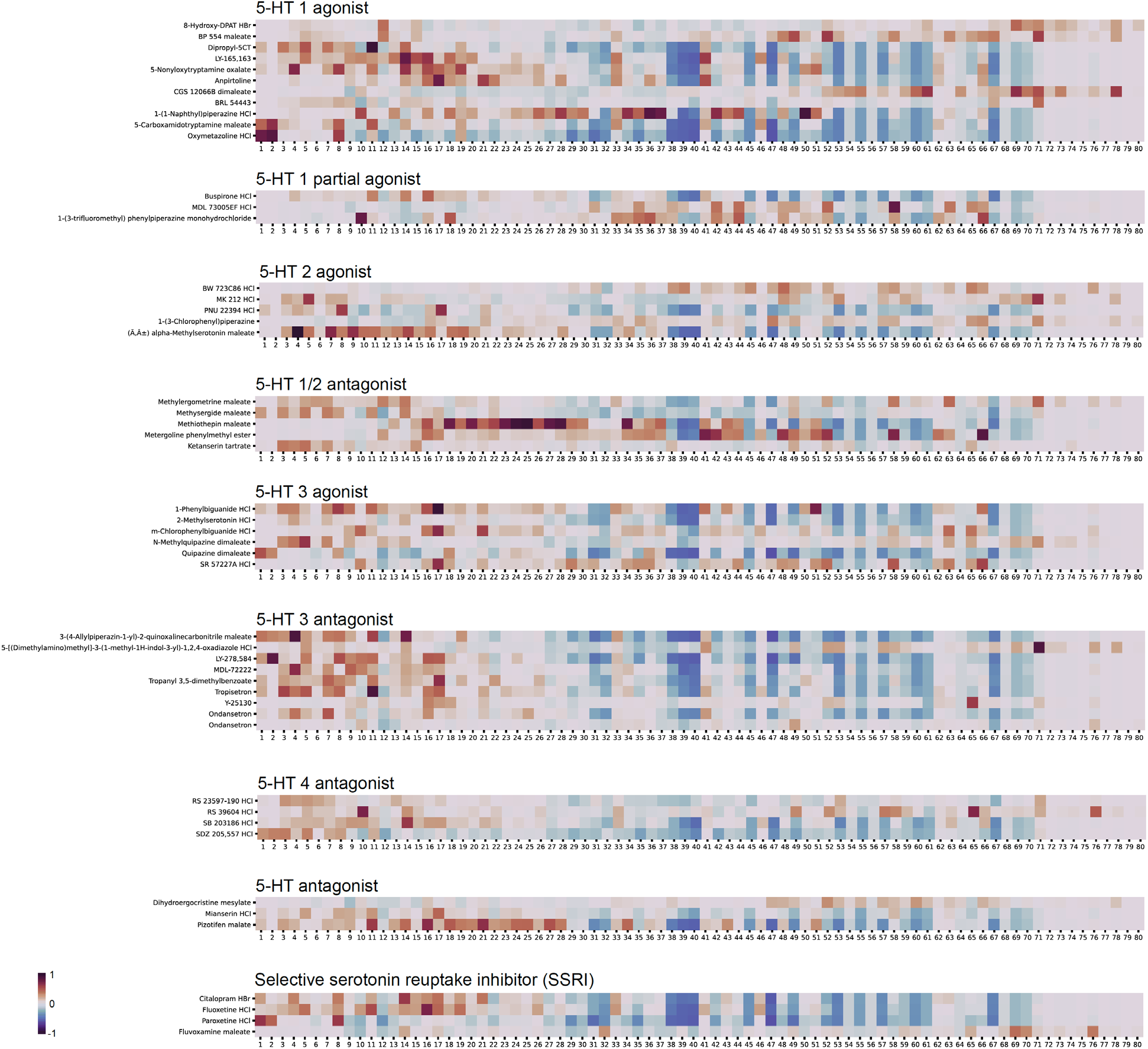
Behavioral fingerprints of serotonin pathway modulators, grouped by drug effects.

**Supplementary Figure 6.**
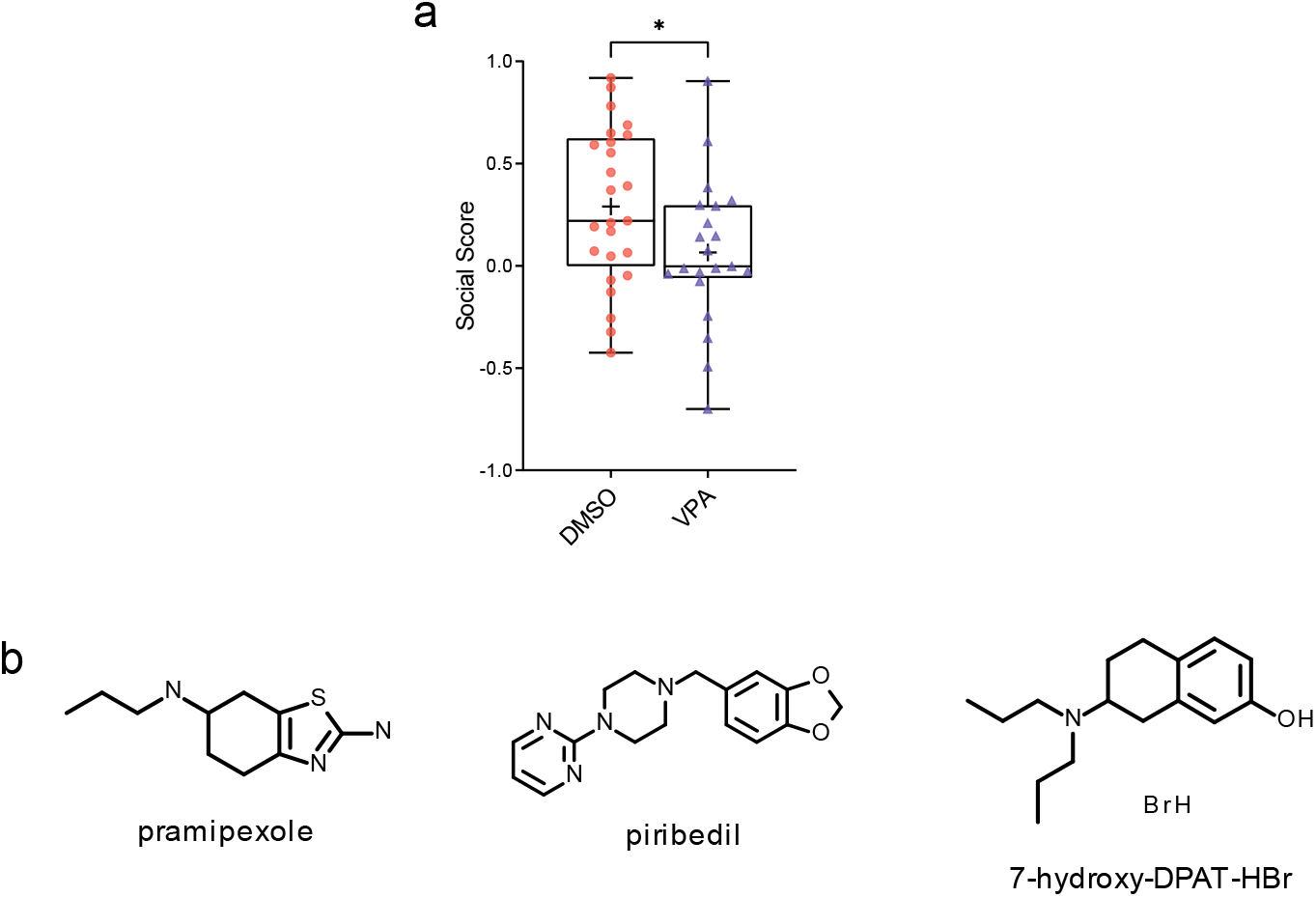
(**a**) Boxplot showing social preference (social score) of fish treated by DMSO (n=25) or valproic acid (VPA; n=21) during the first 3 days of embryonic development. *: *p*<0.05. (**b**) Chemical structures of the D3 agonists pramipexole, piribedil, and 7-hydroxy-DPAT-HBr.

**Supplementary Video 1**. Video recording of a pair of fish interacting in a ZeChat unit. Each unit is divided into two arenas by a transparent window.

**Supplementary Video 2**. Video recording of 40 pairs of fish interacting in a full-sized ZeChat test array.

**Supplementary Video 3**. A combination of 4 processed clips of the same video recording, showing the intermediate and final outcomes of image preprocessing.

**Supplementary Video 4**. Side-by-side view of fish’s behavioral recording and its trajectory on ZeChat map in real-time to visualize how a fish’s behavior translates to datapoint embeddings in the ZeChat map.

**Supplementary Video 5**. Video recordings of wild type (DMSO) and dopamine D3 agonist-treated (10 μM piribedil) fish. Demonstrating a more intense interaction pattern between pairs of D3 agonist-treated fish compared to the wild type.

## REFERENCES

1 Mandell, D. S. et al. Psychotropic medication use among Medicaid-enrolled children with autism spectrum disorders. Pediatrics 121, e441–448, doi:10.1542/peds.2007-0984 (2008).

2 Downs, J. et al. Clinical predictors of antipsychotic use in children and adolescents with autism spectrum disorders: a historical open cohort study using electronic health records. Eur Child Adolesc Psychiatry 25, 649–658, doi:10.1007/s00787-015-0780-7 (2016).

3 Geng, Y. & Peterson, R. T. The zebrafish subcortical social brain as a model for studying social behavior disorders. Dis Model Mech 12, doi:10.1242/dmm.039446 (2019).

4 Kokel, D. et al. Rapid behavior-based identification of neuroactive small molecules in the zebrafish. Nat Chem Biol 6, 231–237, doi:10.1038/nchembio.307 (2010).

5 Bruni, G. et al. Zebrafish behavioral profiling identifies multitarget antipsychotic-like compounds. Nat Chem Biol 12, 559–566, doi:10.1038/nchembio.2097 (2016).

6 Rihel, J. et al. Zebrafish behavioral profiling links drugs to biological targets and rest/wake regulation. Science 327, 348–351, doi:10.1126/science.1183090 (2010).

7 Jordi, J. et al. High-throughput screening for selective appetite modulators: A multibehavioral and translational drug discovery strategy. Sci Adv 4, eaav1966, doi:10.1126/sciadv.aav1966 (2018).

8 Dreosti, E., Lopes, G., Kampff, A. R. & Wilson, S. W. Development of social behavior in young zebrafish. Front Neural Circuits 9, 39, doi:10.3389/fncir.2015.00039 (2015).

9 Stednitz, S. J. et al. Forebrain Control of Behaviorally Driven Social Orienting in Zebrafish. Curr Biol 28, 2445–2451 e2443, doi:10.1016/j.cub.2018.06.016 (2018).

10 Miller, N. & Gerlai, R. Quantification of shoaling behaviour in zebrafish (Danio rerio). Behav Brain Res 184, 157–166, doi:10.1016/j.bbr.2007.07.007 (2007).

11 Tang, W. et al. Genetic Control of Collective Behavior in Zebrafish. iScience 23, 100942, doi:10.1016/j.isci.2020.100942 (2020).

12 Berman, G. J., Choi, D. M., Bialek, W. & Shaevitz, J. W. Mapping the stereotyped behaviour of freely moving fruit flies. J R Soc Interface 11, doi: 10.1098/rsif.2014.0672 (2014).

13 Farneback, G. Two-frame motion estimation based on polynomial expansion. Lect Notes Comput Sc 2749, 363–370, doi:DOI 10.1007/3-540-45103-x_50 (2003).

14 Hinton, L. J. P. v. d. M. a. G. E. Visualizing High-Dimensional Data Using t-SNE. Journal of Machine Learning Research 9, 2579–2605 (2008).

15 Andrej Gisbrecht, A. S., Barbara Hammer. Parametric nonlinear dimensionality reduction using kernel t-SNE. Neurocomputing 147, 71–82 (2015).

16 Gunaydin, L. A. & Deisseroth, K. Dopaminergic Dynamics Contributing to Social Behavior. Cold Spring Harb Symp Quant Biol 79, 221–227, doi:10.1101/sqb.2014.79.024711 (2014).

17 Kiser, D., Steemers, B., Branchi, I. & Homberg, J. R. The reciprocal interaction between serotonin and social behaviour. Neurosci Biobehav Rev 36, 786–798, doi:10.1016/j.neubiorev.2011.12.009 (2012).

18 Pellissier, L. P., Gandia, J., Laboute, T., Becker, J. A. J. & Le Merrer, J. mu opioid receptor, social behaviour and autism spectrum disorder: reward matters. Br J Pharmacol 175, 2750–2769, doi:10.1111/bph.13808 (2018).

19 Nicolini, C. & Fahnestock, M. The valproic acid-induced rodent model of autism. Exp Neurol 299, 217–227, doi:10.1016/j.expneurol.2017.04.017 (2018).

20 Chen, J. et al. Developmental and behavioral alterations in zebrafish embryonically exposed to valproic acid (VPA): An aquatic model for autism. Neurotoxicol Teratol 66, 8–16, doi:10.1016/j.ntt.2018.01.002 (2018).

21 Geng, Y. et al. Top2a promotes the development of social behavior via PRC2 and H3K27me3. bioRxiv, doi:10.1101/2021.09.20.461107 (2021).

22 Branson, K., Robie, A. A., Bender, J., Perona, P. & Dickinson, M. H. High-throughput ethomics in large groups of Drosophila. Nat Methods 6, 451–457, doi:10.1038/nmeth.1328 (2009).

23 Dankert, H., Wang, L., Hoopfer, E. D., Anderson, D. J. & Perona, P. Automated monitoring and analysis of social behavior in Drosophila. Nat Methods 6, 297–303, doi: 10.1038/nmeth.1310 (2009).

24 Laan, A., Iglesias-Julios, M. & de Polavieja, G. G. Zebrafish aggression on the sub-second time scale: evidence for mutual motor coordination and multi-functional attack manoeuvres. R Soc Open Sci 5, 180679, doi:10.1098/rsos.180679 (2018).

25 Hong, W. et al. Automated measurement of mouse social behaviors using depth sensing, video tracking, and machine learning. Proc Natl Acad Sci U S A 112, E5351–5360, doi:10.1073/pnas.1515982112 (2015).

26 de Chaumont, F. et al. Computerized video analysis of social interactions in mice. Nat Methods 9, 410–417, doi:10.1038/nmeth.1924 (2012).

27 Harpaz, R., Tkacik, G. & Schneidman, E. Discrete modes of social information processing predict individual behavior of fish in a group. Proc Natl Acad Sci U S A 114, 10149–10154, doi:10.1073/pnas.1703817114 (2017).

28 Brown, A. E., Yemini, E. I., Grundy, L. J., Jucikas, T. & Schafer, W. R. A dictionary of behavioral motifs reveals clusters of genes affecting Caenorhabditis elegans locomotion. Proc Natl Acad Sci U S A 110, 791–796, doi:10.1073/pnas.1211447110 (2013).

29 Stephens, G. J., Bueno de Mesquita, M., Ryu, W. S. & Bialek, W. Emergence of long timescales and stereotyped behaviors in Caenorhabditis elegans. Proc Natl Acad Sci U S A 108, 7286–7289, doi:10.1073/pnas.1007868108 (2011).

30 Costa, A. C., Ahamed, T. & Stephens, G. J. Adaptive, locally linear models of complex dynamics. Proc Natl Acad Sci U S A 116, 1501–1510, doi:10.1073/pnas.1813476116 (2019).

31 Stephens, G. J., Johnson-Kerner, B., Bialek, W. & Ryu, W. S. Dimensionality and dynamics in the behavior of C. elegans. PLoS Comput Biol 4, e1000028, doi: 10.1371/journal.pcbi.1000028 (2008).

32 Broekmans, O. D., Rodgers, J. B., Ryu, W. S. & Stephens, G. J. Resolving coiled shapes reveals new reorientation behaviors in C. elegans. Elife 5, doi: 10.7554/eLife.17227 (2016).

33 Gomez-Marin, A., Stephens, G. J. & Brown, A. E. Hierarchical compression of Caenorhabditis elegans locomotion reveals phenotypic differences in the organization of behaviour. J R Soc Interface 13, doi:10.1098/rsif.2016.0466 (2016).

34 Szigeti, B., Deogade, A. & Webb, B. Searching for motifs in the behaviour of larval Drosophila melanogaster and Caenorhabditis elegans reveals continuity between behavioural states. J R Soc Interface 12, 20150899, doi:10.1098/rsif.2015.0899 (2015).

35 Berman, G. J., Bialek, W. & Shaevitz, J. W. Predictability and hierarchy in Drosophila behavior. Proc Natl Acad Sci U S A 113, 11943–11948, doi:10.1073/pnas.1607601113 (2016).

36 Todd, J. G., Kain, J. S. & de Bivort, B. L. Systematic exploration of unsupervised methods for mapping behavior. Phys Biol 14, 015002, doi: 10.1088/1478-3975/14/1/015002 (2017).

37 DeAngelis, B. D., Zavatone-Veth, J. A. & Clark, D. A. The manifold structure of limb coordination in walking Drosophila. Elife 8, doi:10.7554/eLife.46409 (2019).

38 Pereira, T. D. et al. Fast animal pose estimation using deep neural networks. Nat Methods 16, 117–125, doi:10.1038/s41592-018-0234-5 (2019).

39 Cande, J. et al. Optogenetic dissection of descending behavioral control in Drosophila. Elife 7, doi:10.7554/eLife.34275 (2018).

40 Marques, J. C., Lackner, S., Felix, R. & Orger, M. B. Structure of the Zebrafish Locomotor Repertoire Revealed with Unsupervised Behavioral Clustering. Curr Biol 28, 181–195 e185, doi:10.1016/j.cub.2017.12.002 (2018).

41 Mearns, D. S., Donovan, J. C., Fernandes, A. M., Semmelhack, J. L. & Baier, H. Deconstructing Hunting Behavior Reveals a Tightly Coupled Stimulus-Response Loop. Curr Biol 30, 54–69 e59, doi:10.1016/j.cub.2019.11.022 (2020).

42 Semmelhack, J. L. et al. A dedicated visual pathway for prey detection in larval zebrafish. Elife 3, doi:10.7554/eLife.04878 (2014).

43 Wiltschko, A. B. et al. Mapping Sub-Second Structure in Mouse Behavior. Neuron 88, 1121–1135, doi:10.1016/j.neuron.2015.11.031 (2015).

44 Markowitz, J. E. et al. The Striatum Organizes 3D Behavior via Moment-to-Moment Action Selection. Cell 174, 44–58 e17, doi:10.1016/j.cell.2018.04.019 (2018).

45 Klibaite, U., Berman, G. J., Cande, J., Stern, D. L. & Shaevitz, J. W. An unsupervised method for quantifying the behavior of paired animals. Phys Biol 14, 015006, doi: 10.1088/1478-3975/aa5c50 (2017).

46 Klibaite, U. & Shaevitz, J. W. Paired fruit flies synchronize behavior: Uncovering social interactions in Drosophila melanogaster. PLoS Comput Biol 16, e1008230, doi:10.1371/journal.pcbi.1008230 (2020).

47 Romero-Ferrero, F., Bergomi, M. G., Hinz, R. C., Heras, F. J. H. & de Polavieja, G. G. idtracker.ai: tracking all individuals in small or large collectives of unmarked animals. Nat Methods, doi:10.1038/s41592-018-0295-5 (2019).

48 Shuo Hong Wang, J. Z., Xiang Liu, Zhi-Ming Qian, Ye Liu, Yan Qiu Chen. 3D tracking swimming fish school with learned kinematic model using LSTM network. IEEE International Conference on Acoustics, Speech and Signal Processing (ICASSP), doi:10.1109/ICASSP.2017.7952320 (2017).

49 Heras, F. J. H., Romero-Ferrero, F., Hinz, R. C. & de Polavieja, G. G. Deep attention networks reveal the rules of collective motion in zebrafish. PLoS Comput Biol 15, e1007354, doi:10.1371/journal.pcbi.1007354 (2019).

50 Hood, S. D. et al. Dopaminergic challenges in social anxiety disorder: evidence for dopamine D3 desensitisation following successful treatment with serotonergic antidepressants. J Psychopharmacol 24, 709–716, doi:10.1177/0269881108098144 (2010).

51 Kagaya, T. et al. Dopamine D3 agonists disrupt social behavior in rats. Brain Res 721, 229–232, doi:10.1016/0006-8993(96)00288-0 (1996).

52 Gendreau, P. L., Petitto, J. M., Petrova, A., Gariepy, J. & Lewis, M. H. D(3) and D(2) dopamine receptor agonists differentially modulate isolation-induced social-emotional reactivity in mice. Behav Brain Res 114, 107–117, doi:10.1016/s0166-4328(00)00193-5 (2000).

53 Hoffman, E. J. et al. Estrogens Suppress a Behavioral Phenotype in Zebrafish Mutants of the Autism Risk Gene, CNTNAP2. Neuron 89, 725–733, doi:10.1016/j.neuron.2015.12.039 (2016).

54 Mwaffo, V., Zhang, P., Romero Cruz, S. & Porfiri, M. Zebrafish swimming in the flow: a particle image velocimetry study. PeerJ 5, e4041, doi: 10.7717/peerj.4041 (2017).

55 Lopes, G. et al. Bonsai: an event-based framework for processing and controlling data streams. Front Neuroinform 9, 7, doi:10.3389/fninf.2015.00007 (2015).

56 Zivkovic, Z. & van der Heijden, F. Efficient adaptive density estimation per image pixel for the task of background subtraction. Pattern Recogn Lett 27, 773–780, doi: 10.1016/j.patrec.2005.11.005 (2006).

